# Highly potent photoinactivation of bacteria using a water soluble, cell permeable, DNA-binding photosensitizer

**DOI:** 10.1101/2021.06.11.448055

**Authors:** Elyse M. Digby, Tianyi Ma, Joshua N. Milstein, Andrew A. Beharry

## Abstract

Antimicrobial photodynamic therapy (APDT) employs a photosensitizer, light, and molecular oxygen to treat infectious diseases via oxidative damage, with a low likelihood for the development of resistance. For optimal APDT efficacy, photosensitizers with cationic charges that can permeate bacteria cells and bind intracellular targets are desired to not limit oxidative damage to the outer bacterial structure. Here we report the application of brominated DAPI (Br-DAPI), a water-soluble, DNA-binding photosensitizer for eradication of both gram-negative and gram-positive bacteria (as demonstrated on N99 *E. coli* and *B. subtilis*, respectively). We observe intracellular uptake of Br-DAPI, ROS-mediated bacterial cell death via 1- and 2-photon excitation, and selective photocytotoxicity of bacteria over mammalian cells. Photocytotoxicity of both N99 *E. coli* and *B. subtilis* occurred at sub-micromolar concentrations (IC_50_ = 0.2 μM – 0.4 μM) and low light doses (5-minute irradiation times, 4.5 J cm^−2^ dose) making it superior to commonly employed APDT phenothiazinium photosensitizers such as methylene blue. Given its high potency and 2-photon excitability, Br-DAPI is a promising novel photosensitizer for in vivo APDT applications.

## INTRODUCTION

The World Health Organization classifies antimicrobial resistance as one of the top ten global health and development threats^1^. Due to the overuse of antibiotics, there exist an alarming number of antibiotic resistant bacteria, which makes effective treatment and inactivation of these pathogens challenging^2^. To combat this global crisis, alternative treatments that kill bacteria without the risk of inducing resistance are highly desirable. Over the years, some alternatives have included phage therapy, antibody treatment, and delivery of antimicrobial peptides^3^; however, these therapies suffer from immunogenicity, high cost of production, and susceptibility to proteolytic degradation, respectively^3^. Moreover, in contrast to conventional antibiotics, most of the alternative approaches (e.g., phage therapy, CRISPR/Cas9, antibodies) are strain-specific resulting in a need for different therapies to treat different types of infections.

An attractive alternative to antibiotics is antimicrobial photodynamic therapy (APDT). Although discovered over 100 years ago, its use has gained renewed interest as antibiotics have begun to fail against drug resistant bacteria^4^. APDT requires three main components – a photosensitizer (PS), wavelength-specific light, and molecular oxygen. Upon irradiation, the PS can produce reactive oxygen species (ROS)^5^, which can oxidatively damage surrounding biomolecules such as lipids, proteins and nucleic acids, ultimately leading to cell death^5^. The non-selective, multi-target, ROS-induced damage renders resistance development highly improbable^6–8^ and enables broad spectrum activity against bacteria^4^. Despite its advantages over other antimicrobial therapies, the use of APDT for treating bacterial infections has been limited by the requirement for high concentrations of the PS and high doses of light to completely eliminate infectious bacteria cells, leading to surrounding host cell damage. Since most PSs have poor solubility in water, high concentrations often result in aggregation, which leads to poor APDT efficacy^9, 10^. To overcome this issue, PSs may be combined with other agents such as antimicrobial peptides^11^, liposomal systems^12^ or biodegradable polymers^13^, or have small cationic groups or other moieties^14–16^ conjugated to the PS. An enhanced permeability improves intracellular ROS production thereby reducing concentrations and light doses needed to achieve high efficacy against bacteria, all while avoiding host tissue damage.^17^ Although positive results have been attained with combinatorial approaches, a PS applied as a single agent capable of exerting high potency at low concentrations and light doses in both gram-negative and gram-positive bacteria would be highly desirable for APDT.

We previously developed a DNA-targeting PS derived from the DNA binding/nucleic acid stain 4’,6-diamidino-2-phenylindole (DAPI)^18^. We found brominated DAPI (Br-DAPI) could bind DNA and fluoresce like native DAPI, but had an additional property of producing ROS upon irradiation. Due to the strong DNA binding of Br-DAPI and the short diffusion distance of ROS, irradiation caused double-strand DNA breaks that led to cancer cell death^18^. We hypothesized that Br-DAPI’s low molecular weight (355 Da) and poly-cationic charge (2+) would permit high permeation in bacteria and given its strong DNA binding affinity would enable direct ROS-induced damage towards DNA rendering it an effective, stand-alone agent for APDT.

## RESULTS AND DISCUSSION

Br-DAPI (Figure 1) was synthesized starting from commercially available DAPI dihydrochloride treated with 3 equiv. of *N*-bromosuccinimide and purified by RP-HPLC as previously described^18^. An extinction coefficient of 33 000 (± 533) M^−1^ cm^−1^ was measured by quantitative H-NMR (see Experimental and Figure S1) and used for concentration measurements.

**Figure 1.**
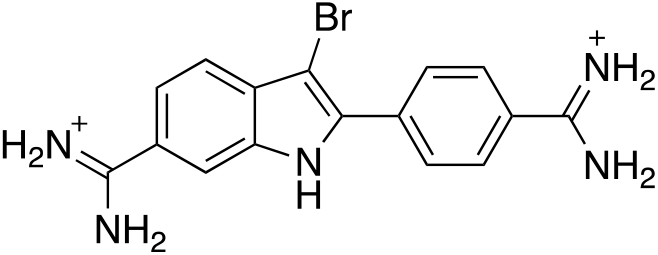
Structure of the DNA binding photosensitizer, Br-DAPI.

We first investigated the intracellular uptake of Br-DAPI in bacteria. It is known that mono-cationic PSs can penetrate deeply through the single-layered cell wall/membrane of gram-positive bacteria, while poly-cationic PSs are usually required for penetration in gram-negative bacteria due to their bilayer membrane structure^4, 17, 19^. Since Br-DAPI is inherently poly-cationic at physiological pH, we reasoned it would display strong electrostatic binding to the negatively charged lipopolysaccharides in both gram-positive and gram-negative bacteria. To test this, we selected the gram-negative bacterial strain N99 *E. coli*, and the gram-positive bacterial strain *B. subtilis*. Since Br-DAPI exhibits low fluorescence in solution (Φ_F_ = 0.005) and increases in cyan fluorescence when bound to DNA^18^, we can directly assess intracellular Br-DAPI uptake, or more specifically, DNA binding, using fluorescence microscopy. We incubated bacteria with Br-DAPI (0.83 μM) in LB broth and, without washing, directly imaged the immobilized bacteria sandwiched between an agarose pad and the coverslip. Within 5 minutes, we observed diffuse cyan fluorescent signals in both the N99 *E. Coli* and *B. subtilis* above background, with similar mean fluorescence intensities (Figure 2 and Figure S2). This suggests Br-DAPI is capable of rapid intracellular entry with similar efficiencies in both gram-negative and gram-positive bacteria, and given the enhanced fluorescence response, at least some fraction is binding to DNA.

**Figure 2.**
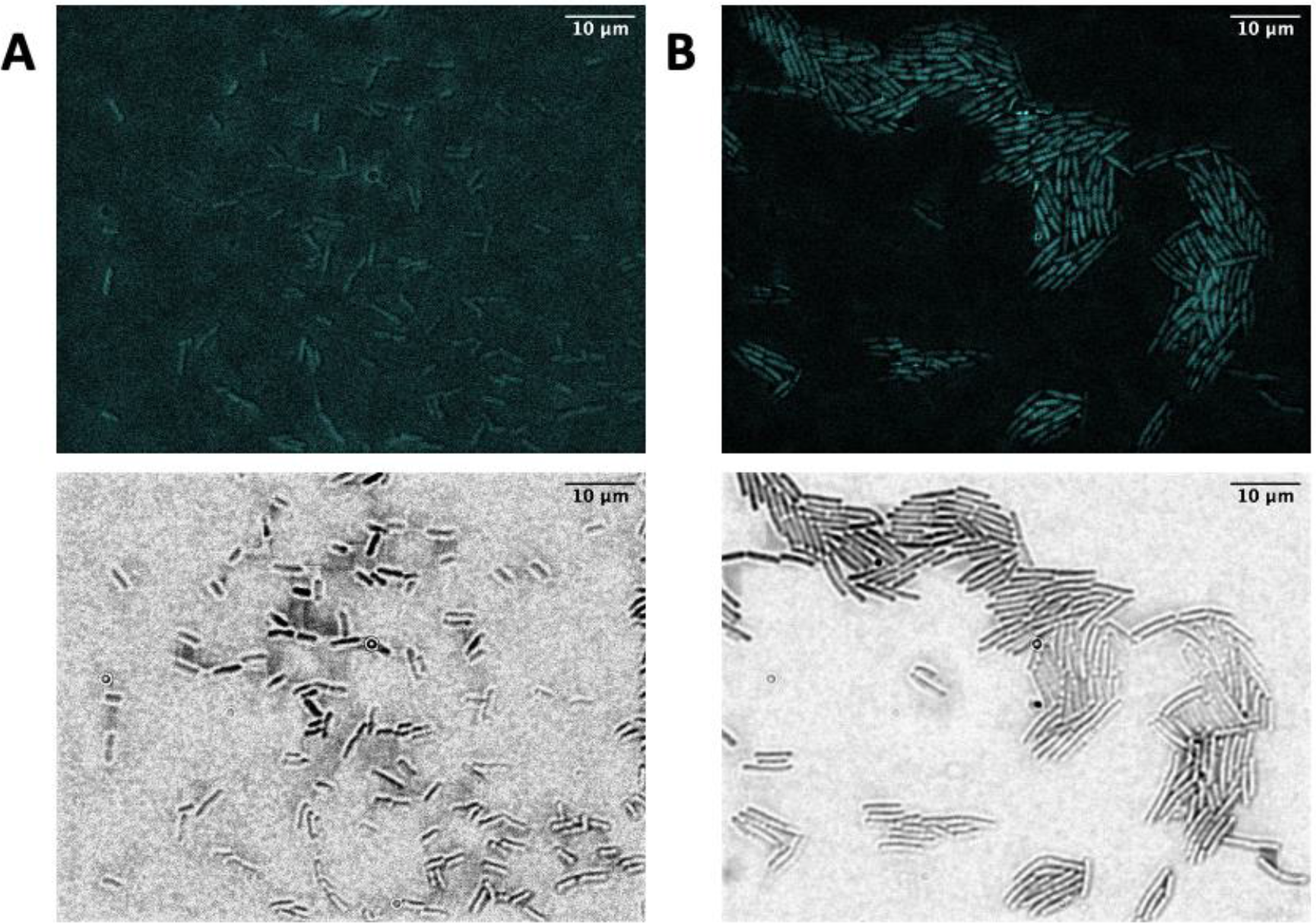
(A) N99 *E. coli* and (B) *B. subtilis* incubated with Br-DAPI (0.83 μM) for 5 minutes. Cyan fluorescence in bacteria above background (Figure S2 (C, D)) demonstrates intracellular uptake. 100x magnification, scale bar = 10 μm. Imaged using DAPI filter cube: λ_ex_ = 387 nm, λ_em_ = 447 nm bandpass filter.

To determine whether intracellular Br-DAPI would produce ROS upon irradiation, we used the general ROS sensor, 2’,7’-dichlorofluorescin diacetate (DCFH_2_-DA). DCFH_2_-DA is cell permeable and non-fluorescent, but can be converted to a green fluorescent product, DCF, upon oxidation via ROS^20, 21^. Standardized suspensions of N99 *E. coli* and *B. subtilis* (OD_600 nm_ = 0.10) were incubated with DCFH_2_-DA (10 μM) and Br-DAPI (0.25 μM) for 5 minutes. Bacteria were then irradiated with a 365 nm LED (15 mW cm^−2^) for 1 minute for a total radiance dose of 0.9 J cm^−2^. Measuring the fluorescence of DCF, we found Br-DAPI produced approximately 11-fold and 4-fold more ROS upon irradiation in N99 *E.* coli and *B.* subtilis, respectively, compared to non-irradiated (dark) controls (Figure 3). Given Br-DAPI’s ability to permeate bacteria cells, bind DNA, and produce ROS, we asked whether a high degree of photocytotoxicity can be achieved.

**Figure 3.**
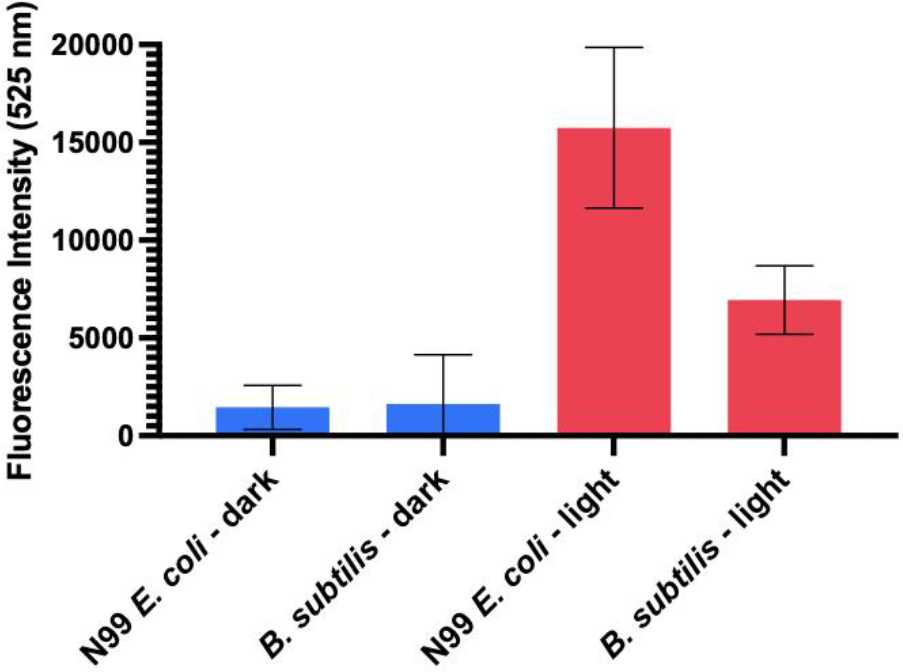
Standardized suspensions of N99 *E. coli* and *B. subtilis* were incubated with DCFH_2_-DA (10 μM) and Br-DAPI (0.25 μM). Bacteria were treated with or without 0.9 J cm^−2^ of irradiation (365 nm LED). ROS production was monitored by the fluorescence of the general ROS sensor DCFH_2_-DA, which produces the green fluorescent product DCF upon oxidation.

Standardized suspensions of N99 *E. coli* and *B. subtilis* were incubated with Br-DAPI (0 μM - 0.83 μM) and irradiated for 5 minutes with a 365 nm LED (15 mW cm^−2^, 4.5 J cm^−2^). Under these irradiation conditions, untreated bacteria remain viable (Figure S3). Bacteria were then diluted and grown overnight on agar and assayed for colony forming units (CFU) 24 hours post treatment. Br-DAPI induced a significant decrease in CFU in a dose-dependent manner under light (IC_50_ = 0.20 μM for N99 *E. Coli*, and IC_50_ = 0.42 μM for *B. subtilis*) (Figure 4). At 1 μM, nearly all colonies were killed under light in both N99 *E. Coli* and *B. subtilis*, with no dark toxicity (Figure 4). The 2-fold higher IC_50_ for *B. subtilis* is consistent with the ~2x less ROS produced compared to N99 *E. coli*. Performing the same experiment with native DAPI, which contains no photosensitizing properties, led to no photocytotoxicity (Figure S4), thereby confirming that Br-DAPI induces toxicity in gram-negative and gram-positive bacteria via light-induced ROS production.

**Figure 4.**
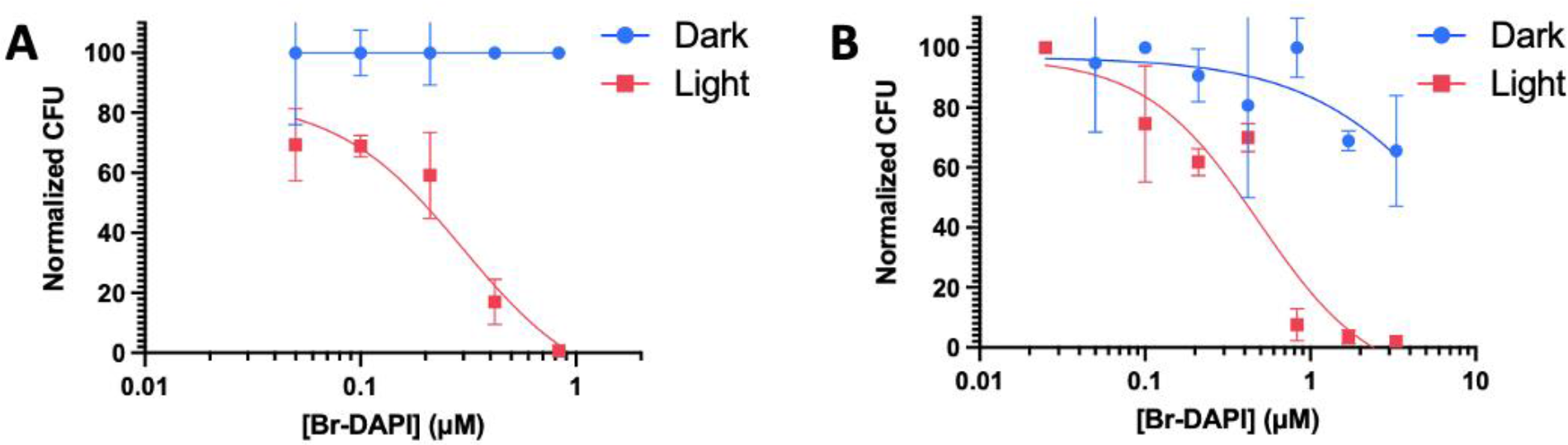
Standardized suspensions of A) N99 *E. coli* and B) *B. subtilis* incubated with different concentrations of Br-DAPI, treated with or without 365 nm irradiation (4.5 J cm^−2^) then incubated overnight in the dark. CFU were assayed 24 hours post-treatment and normalized to 100% relative to untreated bacteria (i.e., 0 μM compound without light). N99 *E. coli*: Light IC_50_ = 0.20 μM, Dark IC_50_ > 0.83 μM; *B. subtilis* Light IC_50_ = 0.42 μM Dark IC_50_ >4 μM.

Given that Br-DAPI binds DNA, produces ROS and kills bacteria, we asked whether light-mediated ROS production causes DNA damage. To determine this, we performed a DNA photocleavage experiment as previously described^10^. Bacteria were treated with Br-DAPI (1 μM) for 5 minutes, irradiated (365 nm, 4.5 J cm^−2^) or kept in the dark, then the DNA was extracted and analyzed by agarose gel electrophoresis. Untreated bacteria, irradiated only or treated with Br-DAPI but kept in the dark, showed intact genomic DNA as visualized by SYBR Safe DNA gel stain (Figure 5). In contrast, bacteria treated with Br-DAPI and light reduced total genomic DNA of N99 *E. Coli* and *B. subtilis* by 74% and 73%, respectively (Figure 5). The lack of appearance of new bands at lower molecular weights is assumed to be due to ROS induced double-stranded DNA breaks, which lead to fragments at concentrations too low to detect by staining, as observed in previously reported PS photocleavage experiments^10, 22^. Overall, these experiments confirm that at least one molecular target of Br-DAPI is DNA, which may be one contributor to the observed high potency in gram-negative and gram-positive bacteria.

**Figure 5.**
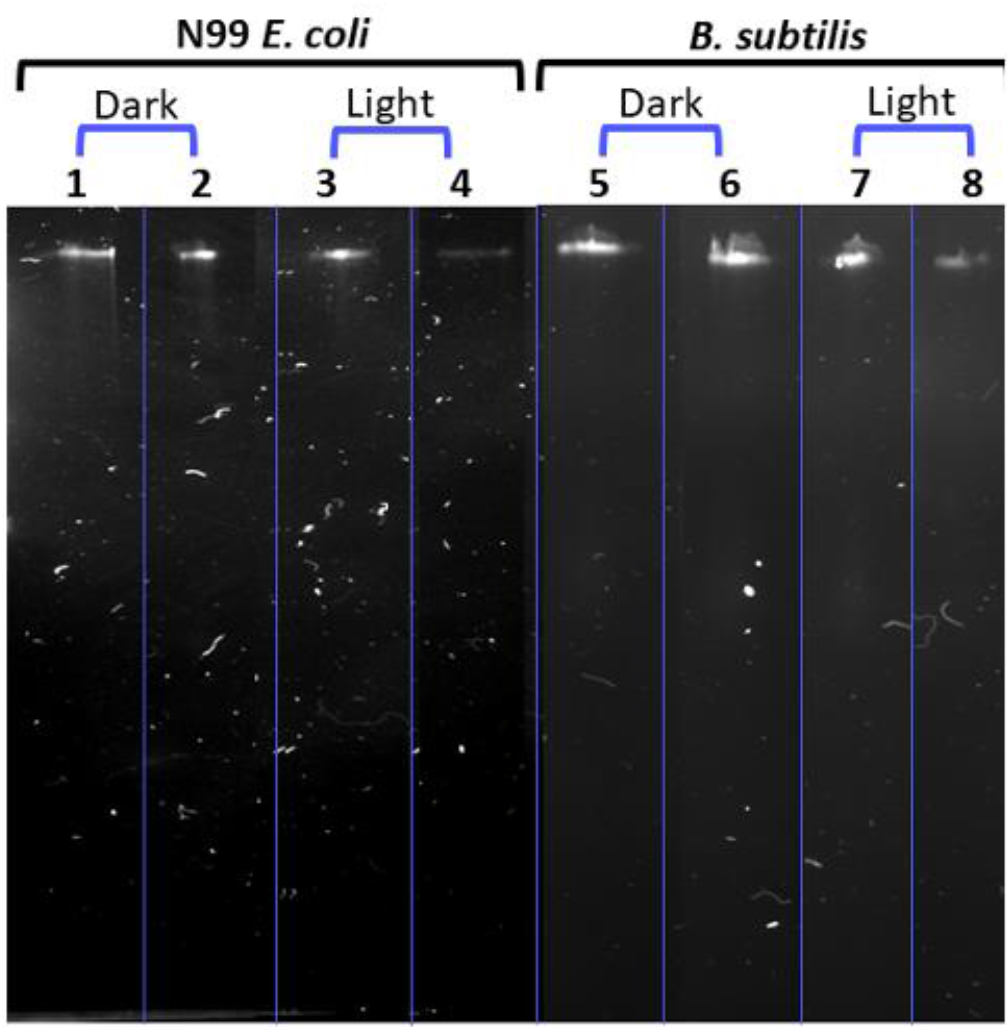
Electrophoretic profile of total DNA extracted from N99 *E. coli* (lanes 1-4) and *B. subtilis* (lanes 5-8) after cells were incubated with Br-DAPI for 5 min in the dark or irradiated with 365 nm LED (4.5 J cm^−2^). Bacteria treated with Br-DAPI and light reduced total genomic DNA content by 74% in N99 *E. coli* and 73% in *B. subtilis*. Lanes 1, 3, 5 and 7 = 0 μM Br-DAPI; Lanes 2, 4, 6 and 8 = 0.5 μM Br-DAPI. DNA was stained with SYBR Safe DNA gel stain.

To compare the degree of photocytotoxicity of Br-DAPI to a commonly employed APDT PS, we performed the same APDT experiments using methylene blue (MB)^23,24^. MB is an FDA approved, amphipathic phenothiazinium salt that exerts its photodynamic efficacy intracellularly and has been traditionally used in a variety of clinical human infections of different etiologies^23,25^. MB was incubated with N99 *E. coli* and *B. subtilis* at the same concentration range (0 μM-0.83 μM) and exposed to the same light dosage (4.5 J cm^−2^) that was used for Br-DAPI, except using a 625 nm LED (11 mW cm^−2^) where MB absorbs maximally. We found that MB did not induce a comparable photocytotoxic effect in either type of bacteria, where CFUs did not reach less than 50% (IC_50_ > 1 μM) (Figure 6). This result is consistent with previous studies of MB requiring higher concentrations and light doses than that used here to exert its photocytotoxic effect^11,23,26–28^. Interestingly, the intracellular ROS levels produced by MB were only ~2-fold less compared to Br-DAPI (Figure 6), despite its IC_50_ (i.e., photocytotoxicity) being at least greater than 5-fold. We hypothesize this may be due to a higher permeability and strong DNA binding ability of Br-DAPI rendering it less susceptible to being pumped out by microbial efflux pumps^25^.

**Figure 6.**
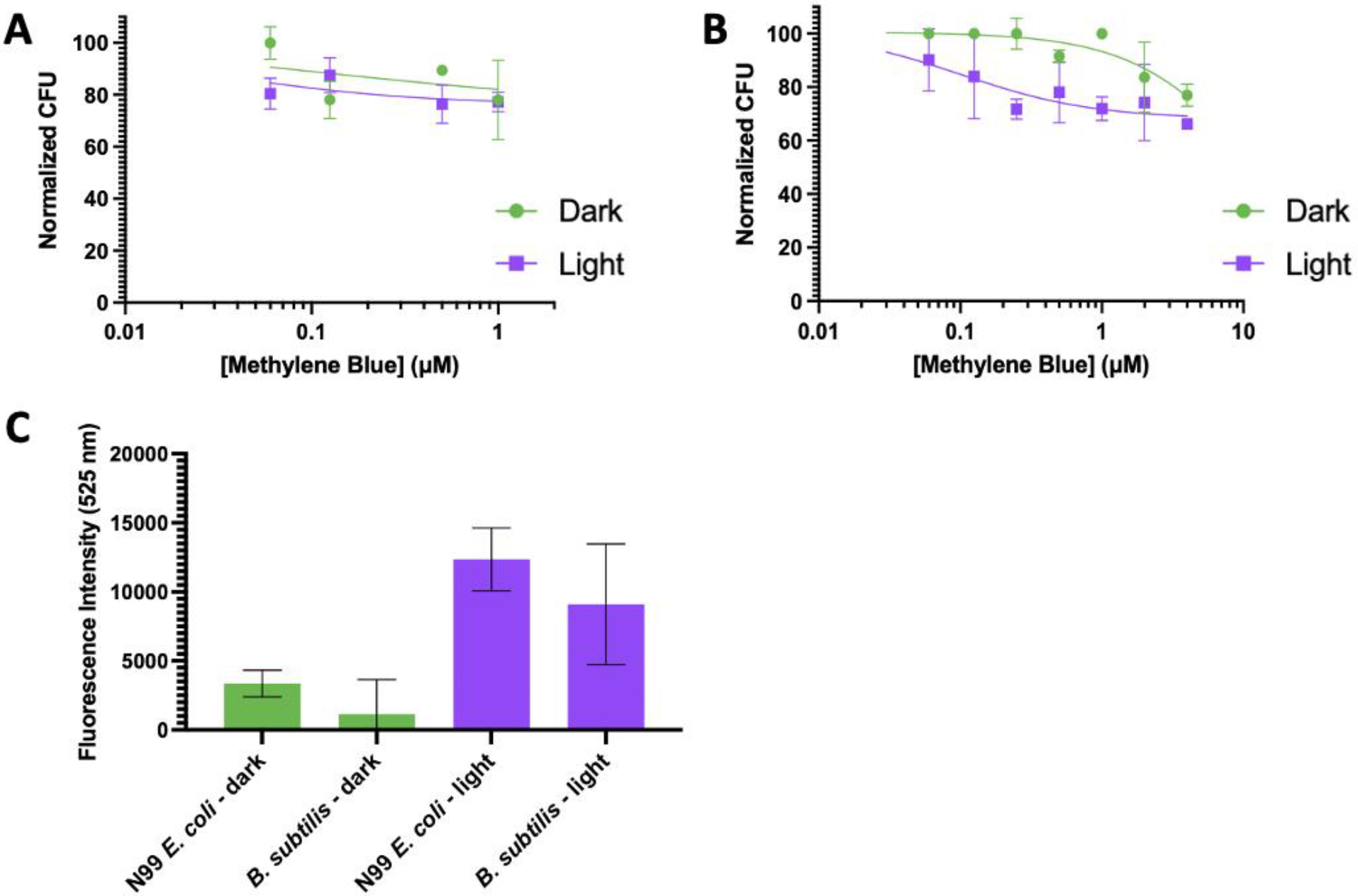
Standardized suspensions of A) N99 *E. coli* and B) *B. subtilis* were incubated with different concentrations of methylene blue, treated with or without 625 nm irradiation (4.5 J cm^−2^), and incubated overnight in the dark. CFU were assayed 24 hours post-treatment and normalized to 100% relative to untreated bacteria (i.e., 0 μM compound without light). C) Standardized suspensions of N99 *E. coli* and *B. subtilis* were incubated with DCFH_2_-DA (10 μM) and methylene blue (0.25 μM). Bacteria were treated with or without 0.9 J cm^−2^ of irradiation with a 625 nm LED. ROS production was monitored by the fluorescence of the general ROS sensor DCFH_2_-DA, which produces the green fluorescent product DCF upon oxidation.

Thus far, a drawback with Br-DAPI is that it requires UV-A excitation, wavelengths that suffer from short tissue penetration depths and background toxicity in vivo. To minimize scattering and absorption of light by biological tissues, longer irradiation wavelengths within a window from 650 nm – 950 nm are desirable to achieve maximal APDT in vivo^29^. Using the notion that native DAPI is 2-photon active^30,31^, we asked whether Br-DAPI is also 2-photon active and can be used for APDT. To explore this, Br-DAPI was incubated with N99 *E. coli* (0 μM −0.83 μM) and irradiated with a 690 nm 2-photon laser (5 minutes, 30 J output power). Under these irradiation conditions, no background photocytotoxicity was observed (Figure S5). We found 2-photon APDT with Br-DAPI led to a dose-dependent photocytotoxic response with a potency similar to that achieved with 1-photon, UV-A irradiation (IC_50_ = 0.30 μM) (Figure 7). These results suggest longer wavelengths of light can be used for Br-DAPI to exert its APDT effect, thereby allowing for deeper tissue penetration and minimizing background phototoxicity in vivo.

**Figure 7.**
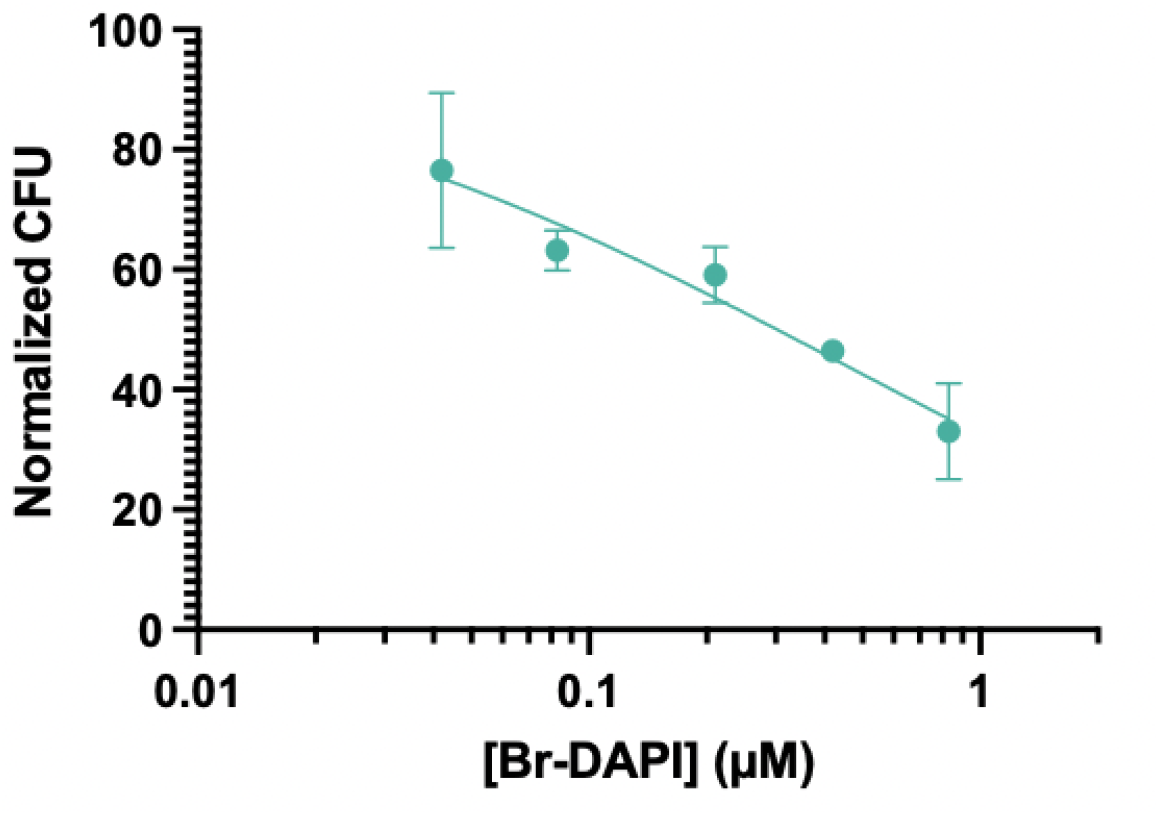
Standardized suspensions of N99 *E. coli* were incubated with different concentrations of Br-DAPI, treated with 690 nm 2-photon laser irradiation (30 J output power), and incubated overnight in the dark. CFU were assayed 24 hours post-treatment and normalized to 100% relative to untreated bacteria (i.e., 0 μM compound without light).

Lastly, the field of targeted APDT aims to kill pathogens without damaging healthy tissue. Cationic PSs are well adapted to APDT as they are not only more efficient at permeating bacteria compared to neutral or negatively charged PS, but they also induce some selectivity over normal cells given the higher density of negative charge on bacteria^32^. However, since mammalian cells are also negatively charged and given the high concentrations required for many clinically relevant PSs to eliminate bacteria (e.g., MB used in concentration range of 0.3 mM - 60 mM)^23^, selectivity for bacteria has been difficult to achieve, leading to the development of targeted strategies by derivatizing PSs^17^. To test whether Br-DAPI has any intrinsic selectivity for bacterial cell death over mammalian cell death, we performed a co-culture APDT experiment using a suspension of N99 *E. coli* with adherent MRC-9 healthy lung cells. Br-DAPI (0.42 μM) was added to the co-culture as well as the cell death stain, propidium iodide (PI). PI is only able to permeate cell membranes that have been damaged, where it can then bind DNA producing red fluorescence^33,34^. Thus, PI fluorescence can be used to distinguish dead cells from live cells. After 5 minutes incubation in the dark, the cells were irradiated for 5 minutes with a 365 nm LED (15 mW cm^−2^, 4.5 J cm^−2^), then incubated in the dark for 30 minutes. We observed PI red fluorescence in only the N99 *E. coli* and not in MRC-9 cells, suggesting only bacteria are dead and not the mammalian cells (Figure 8). This selective killing is consistent with the micromolar concentrations of Br-DAPI required to exert photocytotoxicity on MRC-9 cells only (Figure S6), while only sub-micromolar concentrations are required to kill bacteria (Figure 4 A, B). Moreover, we observed cyan fluorescence from Br-DAPI primarily in the N99 *E. coli* (Figure 4C) suggesting that accumulation in bacteria is higher than that of the MRC-9 at the low micromolar concentration used in the co-culture. Thus, Br-DAPI is capable of inducing selective photocytotoxicity of bacteria in the presence of mammalian cells, making it a promising agent for efficacious APDT in vivo with minimal damage to host cells.

**Figure 8.**
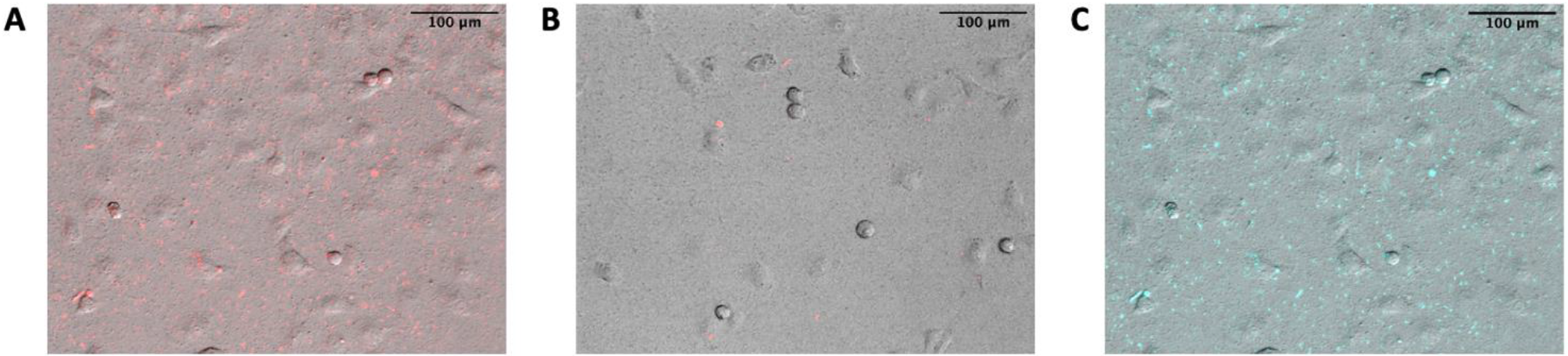
A co-culture of MRC-9 lung cells and a standardized suspension of N99 *E. coli* were incubated with A) Br-DAPI (0.42 μM) and propidium iodide (5 μM), or B) propidium iodide only, treated with 365 nm irradiation (4.5 J cm^−2^). Cells were imaged 30 minutes post-treatment for evidence of cell death which was indicated by the red fluorescence of the propidium iodide cell death fluorescent indicator. (C) The cyan fluorescence from Br-DAPI was selectively observed in bacteria and minimally in mammalian cells. Brightfield images were overlayed with fluorescence of propidium iodide (A, B) or Br-DAPI (C). 16x magnification, scale bar 100 μm. PI imaged using Cy5 filter cube: λ_ex_ = 628 nm, λ_em_ = 692 nm bandpass filter. Br-DAPI imaged using DAPI filter cube: λ_ex_ = 387 nm, λ_em_ = 447 nm bandpass filter.

## CONCLUSION

We have employed our previously developed DNA-binding PS, Br-DAPI, for APDT. The polycationic charge state of Br-DAPI renders the molecule highly water soluble and cell permeable in both gram-negative and gram-positive bacteria despite the different compositions of their cell walls/membranes. The binding of Br-DAPI to DNA allows real-time fluorescence visualization of bacteria, and both intracellular ROS production and complete photoinactivation can be achieved with low light doses (1-photon at 365 nm or 2-photon at 690 nm). Moreover, selective killing of bacteria can be readily achieved in the presence of mammalian cells. Br-DAPI’s sub-micromolar potency in both gram-negative and gram-positive bacteria make it superior to commonly employed phenothiazinium PSs like the FDA approved MB as demonstrated here, and to other recently reported PSs, which in addition to having lower potencies, often photoinactivate either gram-positive or gram-negative bacteria, but not both^10,16,35,36^. Given its high potency, bacterial selectivity over mammalian cells, and 2-photon excitability, Br-DAPI may be a promising stand-alone PS for treating more complex bacterial systems. For example, the treatment of skin infections may benefit from topical applications of Br-DAPI given its large therapeutic window capable of killing bacteria without eukaryotic cell damage, and its ability to photoinactive using long-wavelengths would permit targeting bacteria deep within tissues. Moreover, eradication of bacteria in biofilms are often poor due to inefficient compound penetration and binding to bacteria in the extracellular polymer matrix, resulting in 100-fold higher concentrations of antimicrobial treatments than the determined minimum inhibitory concentrations in culture^37^. Using Br-DAPI, it may be possible to overcome such challenges given its polycationic charge, low molecular weight and already high (sub-micromolar) potency. Such applications are currently being explored.

## METHODS

### Materials and General Methods

All chemicals and instruments were obtained from commercial suppliers. UV-Vis absorption spectra were recorded in a 1.0 cm path length quartz cuvette on a Shimadzu UV-1800 UV-Vis spectrometer. 1-photon irradiation experiments were performed with a mounted 365 nm LED (7.5 nm bandwidth, 360 mW LED Output Power; 8.9 μW mm^−2^ maximum irradiance; M365L2) or a mounted 625 nm LED (17 nm bandwidth, 920 mW LED Output Power; 21.9 μW mm^−2^ maximum irradiance; M625L4) purchased from ThorLabs. 2-photon irradiation experiments were performed on an LSM880 confocal microscope equipped with an InSight®X3™ Tunable Laser. Microscope images were obtained on an Olympus® IX81 Motorized Inverted Research Microscope equipped with a Hamamatsu® C4742-95 CCD camera. Fluorescence imaging was captured using Semrock Brightline Cy5 Fluorescence Filter Cube and Semrock Brightline DAPI Fluorescence Filter Cube. Image processing was performed using ImageJ software.

### Synthesis of Br-DAPI^18^

4’,6-Diamidino-2-phenylindole 2HCl (purchased from Biosynth Carbosynth, 1 eq., 0.0100 g, 0.029 mmol) was dissolved in 1:1 distilled water/acetone (1 mL). Next, *N*-bromosuccinimide (3 eq., 0.0154 g, 0.087 mmol) was added and the reaction mixture was stirred overnight at r.t. protected from light. The crude mixture was dried under reduced pressure and purified directly by RP-HPLC in acetonitrile and 0.1% formic acid in milli-q water (45 min. method with a 25 min. gradient from 5-100% ACN, monitored at 345 nm) – product elutes at 21 min. peak during the gradient. Br-DAPI was obtained as a yellow solid with 45% yield (4.6 mg, 0.013 mmol). HRMS (ESI+) *m/z* calculated for C_16_H_14_BrN_5_ 355.04; found 356.0500. ^1^H NMR (400 MHz, *d_6_*-DMSO) δ 8.46 (s, 2H, 8.13 (d, 8.3 Hz, 2H), 8.01 (s, 1H), 7.98 (d, 4.3 Hz, 2H), 7.67 (d, 8.4 Hz, 1H), 7.56 (d, 8.5 Hz, 1H). ^13^C NMR (101 MHz, *d_6_*-DMSO) δ 168.62, 167.13, 166.07, 137.08, 135.49, 135.28, 131.56, 129.51, 128.73, 123.95, 120.00, 119.56, 113.19, 89.60.

### Br-DAPI Quantitative ^1^H NMR

Br-DAPI was dissolved in *d_6_-*DMSO containing 150 mM 1,4-dioxane as a standard, with a total volume of 400 μL. A ^1^H NMR was run on the sample and analyzed accordingly, where the peak corresponding to 1,4-dioxane integrated to 100.88 ppm. The concentration of Br-DAPI was then determined using Eq. 1 to be 12 mM. The absorbance of Br-DAPI in the NMR sample was measured using a series of dilutions (0-10x). The absorbance at 353 nm was plotted against the sample’s molar concentration to obtain the molar extinction coefficient from the slope of the line using the Beer Lamber equation (Eq. 2). The molar extinction coefficient of Br-DAPI was measured to be 33 000 (±533) M^−1^ cm^−1^ at 353 nm.

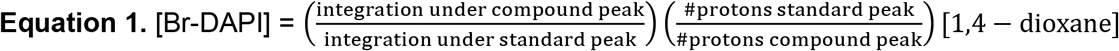

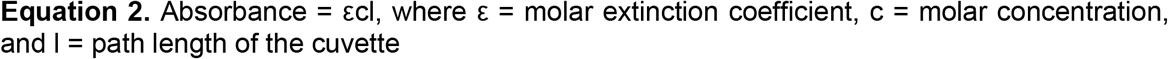

### Concentrations

All stock solutions were prepared in DMSO (Sigma Aldrich). The molar extinction coefficient for all compounds in PBS pH 7.4 were used to measure concentrations of stock solutions – 33 000 (±533) M^−1^ cm^−1^ at 353 nm for Br-DAPI, 27 000 M^−1^ cm^−1^ at 353 nm for DAPI^38^, and 95 000 M^−1^ cm^−1^ at 664 nm for MB^39^.

### Bacterial Strains and Preparation

Colonies of *Escherichia coli* (N99 pXG10sf-GFP) and *Bacillus subtilis* (wild-type) were thawed from frozen concentrated stocks. Bacteria were inoculated from an agar plate into 5 mL of LB broth (ampicillin and tetracycline antibiotics added to N99 *E. coli*) and grown at 37°C overnight with shaking at 250 rpm. The next day, bacteria were regrown by diluting the overnight culture 100x into fresh LB broth (5 mL total volume) (antibiotics added for N99 *E. coli*). Bacteria were grown at 37°C for an additional 2 hours with shaking at 250 rpm until they reached an exponential phase (OD_600_ = 0.10-0.50). Standardized suspensions of bacteria were prepared for all experiments to the same OD_600_.

### Cell Culture

MRC-9 (human lung fibroblasts) were purchased from American Type Culture Collection (ATCC) and cultured in Eagle’s Minimum Essential Medium (EMEM) with sodium pyruvate and L-glutamine (ATCC), supplemented with 10% FBS and 1% antibiotic-antimycotic solution. Cells were cultured in a 75 cm^2^ culture flask at 37°C under 5% CO_2_ in a humidified incubator.

### Fluorescence Imaging for Intracellular Uptake of Br-DAPI

Bacteria (N99 *E. coli* and *B. subtilis*) were cultured overnight and regrown to an OD_600_ = 0.40. Br-DAPI was added to 100 μL suspensions of bacteria for a final concentration of 0.83 μM (<1% DMSO) and incubated at 37°C for 5 minutes before imaging on agarose pads. Agarose pads were prepared by melting 1.5% low-melt agarose in LB broth, placing 1 mL of the mixture onto a micro cover glass (22×22 μm), covering the droplet with a second cover glass, and leaving to solidify at room temperature. After the agarose solidified, the top cover glass was removed, and the agarose was cut into approximately 5×5 mm square pads. 1 μL of bacteria was dropped onto each agar pad and left to air dry for 5 minutes. Agarose pads containing bacteria samples were flipped onto a new cover glass (cells facing onto cover glass), sandwiched between a second cover glass, and sealed into an imaging chamber containing a custom spacer. Br-DAPI’s fluorescence was imaged using a DAPI filter cube (λ_ex_ = 387 nm, λ_em_ = 447 nm bandpass filter) for N99 *E. coli* and *B. subtilis* at 100x magnification, scale bar = 10 μm).

### Detection of ROS Production in Bacteria

Bacteria (N99 *E. coli* and *B. subtilis*) were cultured overnight, regrown, and standardized to an OD_600_ = 0.10. 100 μL standardized suspensions of bacteria, the ROS sensor 2’,7’-dichlorofluorescin diacetate (DCFH_2_DA, 10 μM) and Br-DAPI or MB (0.25, 0.50 μM; < 1% DMSO) were added and incubated at 37°C for 5 minutes in two black 96-well plates (separate plate for dark and light treatments). Bacteria containing Br-DAPI were irradiated with a 365 nm LED for 60 seconds (0.9 J cm^−2^) and bacteria containing MB were irradiated with a 625 nm LED for 84 seconds (0.9 J cm^−2^). The green fluorescence of DCF (oxidized product of the ROS sensor) was measured in bacteria on a multimode microplate reader (Tecan Spark® 20M): λ_ex_=495 nm, λ_em_=525 nm. The fluorescence intensity of DCF for bacteria containing the PSs were corrected for any background fluorescence due to the sensor alone under both dark and light conditions.

### Agarose Gel Electrophoresis of Bacterial Genomic DNA

Genomic DNA was extracted from N99 *E. coli* and *B. subtilis* after treatment as reported in a previous study^10^ using PureLink™ Genomic DNA Mini Kit (purchased from ThermoFisher Sceintific). 1 mL suspensions of bacteria from the overnight culture were incubated with Br-DAPI (0.5 μM; < 1% DMSO) for 5 minutes at 37°C and those designated for light treatment irradiated with a 365 nm LED for 5 minutes (4.5 J cm^−2^). All bacteria samples were then collected by centrifugation at 14 000 *g* for 2 minutes, washed with 0.1% NaCl, and extracted according to the manufacturer’s instructions using 25 μL of the elution buffer per sample. Concentrations of extracted DNA samples were measured using a nano-drop spectrophotometer (NanoPhotometer® P-330) using the ratio of absorbance at 260/280 nm. The extracted DNA was analyzed by gel electrophoresis using 0.8% (w/v) agarose containing SYBR Safe DNA gel stain (5 μL in 5 mL agar solution) in 1X TAE buffer for 45 minutes at 100 V. Gels were imaged using BIORAD ChemiDoc™MP Imaging System.

### Antimicrobial Photocytotoxicity in N99 *E. coli* and *B. subtilis*

#### Optimal light dose determination in N99 *E. coli* for 1-photon PDT

N99 *E. coli* were cultured overnight, regrown, and standardized to an OD_600_ = 0.10. 100 μL of bacteria were added to wells in a 96-well plate and irradiated with a 365 nm LED for times between 0-5 minutes (0-4.5 J cm^−2^).

#### Photosensitizing effect due to bromination of native DAPI in N99 *E. coli*

N99 *E. coli* were cultured overnight, regrown, and standardized to an OD_600_ = 0.10. Br-DAPI or DAPI at increasing concentrations (0-6.70 μM; <1% DMSO) were added to 100 μL of the standardized suspension in two 96-well plates (dark and light) and incubated at 37°C for 5 minutes. Bacteria in the light-designated plate were irradiated with a 365 nm LED for 5 minutes (4.5 J cm^−2^).

#### Comparison of photocytotoxicity due to Br-DAPI and MB in bacteria with 1-photon PDT

Bacteria (N99 *E. coli* and *B. subtilis*) were cultured overnight, regrown, and standardized to an OD_600_ = 0.10. Br-DAPI or MB at increasing concentrations (0 - 0.83 μM for N99 *E. coli*, 0 - 3.3 μM for *B. subtilis*; <1% DMSO) were added to 100 μL of the standardized suspensions in two 96-well plates (dark and light) and incubated at 37°C for 5 minutes. Bacteria in the light-designated plate containing Br-DAPI were irradiated with a 365 nm LED for 5 minutes (4.5 J cm^−2^) and those containing MB were irradiated with a 625 nm LED for 7 minutes (4.5 J cm^−2^).

#### Optimal light dose determination in N99 *E. coli* for 2-photon PDT

N99 *E. coli* were cultured overnight, regrown, and standardized to an OD_600_ = 0.10. 100 μL of bacteria were added to wells in a clear 384-well plate and irradiated with a 690 nm 2-photon laser for 5 minutes with increasing laser power (0-150 J).

#### Photocytotoxicity due to Br-DAPI with 2-photon PDT

N99 *E. coli* were cultured overnight, regrown, and standardized to an OD_600_ = 0.10. Br-DAPI at increasing concentrations (0 - 0.83 μM; <1% DMSO) were added to 100 μL of the standardized suspension in a clear 384-well plate at 37°C for 5 minutes. Bacteria were irradiated with a 690 nm 2-photon laser for 5 minutes (30 J output power).

#### CFU assay for dark and light treated bacteria

6-well cell culture plates (Cell Star, Cat. -No. 657 160) were prepared by melting agarose (purchased from Bio Basic Canada Inc.; AB0015) pouring 3 mL into each well, and leaving to solidify at room temperature. Bacteria from each treatment were diluted in LB broth (1500x N99 *E. coli*, 100x *B. subtilis*), and 5 μL of each were transferred onto 50 μL of LB broth in separate wells. The bacteria mixtures were spread onto the agarose surface with glass beads and left to incubate overnight at 37°C. 24 hours post-treatment, CFU were counted manually – any wells containing <30 or >300 CFU were the lower and upper limit, respectively.

### Photocytotoxicity due to Br-DAPI PDT in co-culture with MRC-9 lung cells and N99 *E. coli*

N99 *E. coli* were cultured overnight and regrown to an OD_600_ = 0.50. MRC-9 lung cells were cultured at a cell density of 40 000 cells well^−1^ in 250 μL of EMEM overnight at 37°C with 5% CO_2_ in an 8-well chamber (ThermoFisher Scientific, Cat. -No. 154534). To a 250 μL suspension of bacteria in LB broth, Br-DAPI for a final concentration of 0.42 μM (<1% DMSO) and propidium iodide (PI, 5 μM) were added. The old media from MRC-9 was removed and the bacteria containing Br-DAPI and PI were added to a well and put to incubate at 37°C for 5 minutes. The co-culture was then irradiated with a 365 nm LED for 5 minutes (4.5 J cm^−2^). The bacteria were transferred to an Eppendorf, washed with 0.1% saline, centrifuged, and then resuspended in DPBS, while the MRC-9 were washed 1x with DPBS. Bacteria were then added back into the well containing MRC-9 and left to incubate at 37°C for 30 minutes for cell death to occur. Cell death was determined by the red fluorescence of PI which will only permeate and bind DNA of dying cells^34,40^. PI and Br-DAPI’s fluorescence were imaged using a Cy5 filter cube (λ_ex_ = 628 nm, λ_em_ = 692 nm bandpass filter) and DAPI (λ_ex_ = 387 nm, λ_em_ = 447 nm bandpass filter) filter cube, respectively. 16x magnification, scale bar = 100 μm.

## ACKNOWLEDGEMENTS

E.M.D and A.A.B. acknowledge support from an NSERC Discovery grant. T. M. and J. N. M. acknowledge support from an NSERC Discovery grant and the New Frontiers in Research Funding Exploration program.

---

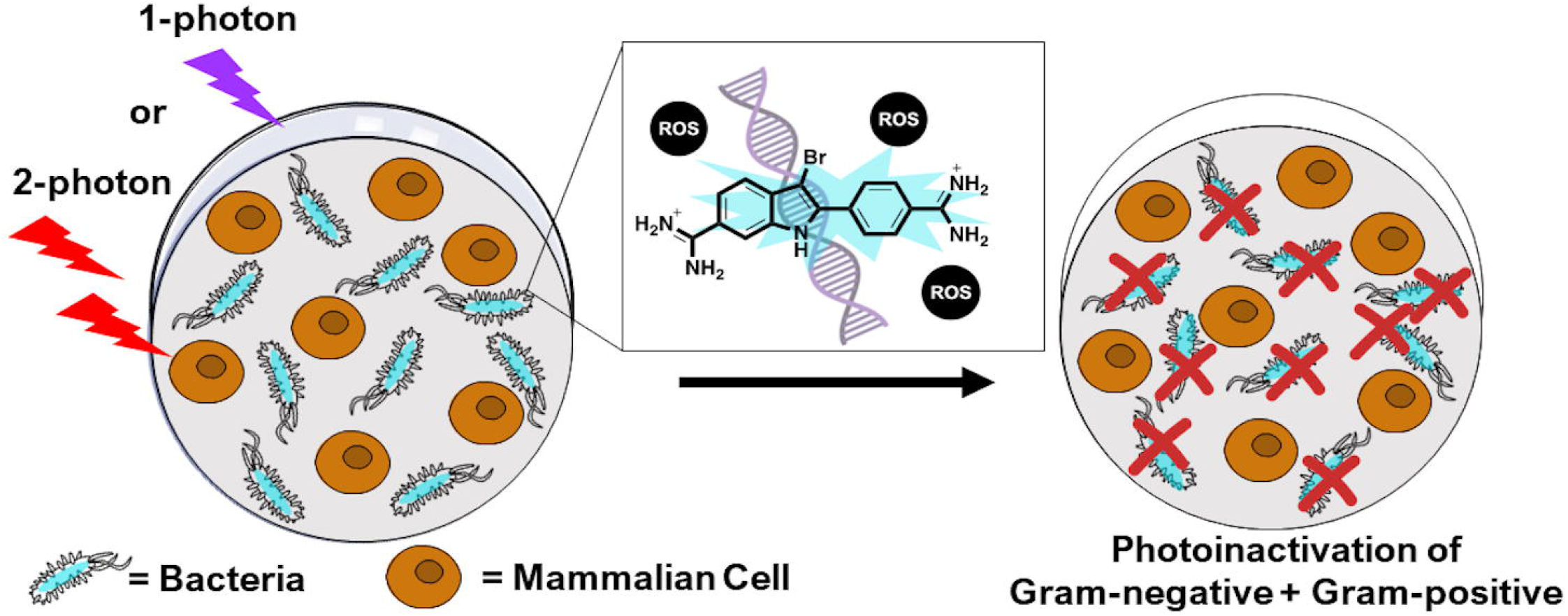

## REFERENCES

1. Global Action Plan on Antimicrobial Resistance; World Health Organization, Ed.

2. Wang, W.; Arshad, M. I.; Khurshid, M.; Rasool, M. H.; Nisar, M. A.; Aslam, M. A.; Qamar, M. U (2018) Antibiotic Resistance : A Rundown of a Global Crisis. Infect. Drug Resist. 11, 1645–1658.

3. Ghosh, C.; Sarkar, P.; Issa, R.; Haldar, J. (2019) Alternatives to Conventional Antibiotics in the Era of Antimicrobial Resistance. Trends Microbiol. 27 (4), 323–338.

4. Jia, Q.; Song, Q.; Li, P.; Huang, W. (2019) Rejuvenated Photodynamic Therapy for Bacterial Infections. Adv. Healthc. Mater. 8 (14), 1–19.

5. Luby, B. M.; Walsh, C. D.; Zheng, G. (2019) Advanced Photosensitizer Activation Strategies for Smarter Photodynamic Therapy Beacons. Angew. Chemie 131 (9), 2580–2591.

6. Maisch, T. (2015) Resistance in Antimicrobial Photodynamic Inactivation of Bacteria. Photochem. Photobiol. Sci. 14, 1518–1526.

7. Wainwright, M.; Maisch, T.; Nonell, S.; Plaetzer, K.; Almeida, A.; Tegos, G. P.; Hamblin, M. R (2017) Photoantimicrobials—Are We Afraid of the Light? Lancet Infect. Dis. 17 (2), 49–55.

8. Cassidy, C. M.; Donnelly, R. F.; Tunney, M. M (2010) Effect of Sub-Lethal Challenge with Photodynamic Antimicrobial Chemotherapy (PACT) on the Antibiotic Susceptibility of Clinical Bacterial Isolates. J. Photochem. Photobiol. B Biol. 99 (1), 62–66.

9. Almeida-Marrero, V.; van de Winckel, E.; Anaya-Plaza, E.; Torres, T.; de la Escosura, A. (2018) Porphyrinoid Biohybrid Materials as an Emerging Toolbox for Biomedical Light Management. Chem. Soc. Rev. 47, 7369–7400.

10. Yang, Z.; Qiao, Y.; Li, J.; Wu, F.; Lin, F. (2020) Novel Type of Water-Soluble Photosensitizer from Trichoderma Reesei for Photodynamic Inactivation of Gram-Positive Bacteria. Langmuir 36 (44), 13227–13235.

11. De Freitas, L. M.; Lorenzón, E. N.; Santos-Filho, N. A.; Zago, L. H. D. P.; Uliana, M. P.; De Oliveira, K. T.; Cilli, E. M.; Fontana, C. R (2018) Antimicrobial Photodynamic Therapy Enhanced by the Peptide Aurein. Sci. Rep. 8 (1), 1–15.

12. Boccalini, G.; Conti, L.; Montis, C.; Bani, D.; Bencini, A.; Berti, D.; Giorgi, C. (2017) Methylene Blue-Containing Liposomes as New Photodynamic Anti-Bacterial Agents. J. Mater. Chem. B 5, 2788–2797.

13. Perni, S.; Preedy, E. C.; Prokopovich, P. (2021) Amplify Antimicrobial Photo Dynamic Therapy Efficacy with Poly - Beta - Amino Esters (PBAEs). Sci. Rep. 11, 1–15.

14. Zhou, J.; Qi, G.; Wang, H. (2016) A Purpurin-Peptide Derivative for Selective Killing of Gram-Positive Bacteria via Insertion into Cell. J. Mater. Chem. B 4, 4855–4861.

15. Zhao, Y.; Ying, J.; Sun, Q.; Ke, M.; Zheng, B.; Huang, J. (2020) A Novel Silicon (IV) Phthalocyanine-Oligopeptide Conjugate as a Highly Efficient Photosensitizer for Photodynamic Antimicrobial Therapy. Dye. Pigment. 172, 107834.

16. Ucuncu, M.; Mills, B.; Duncan, S.; Staderini, M.; Dhaliwal, K.; Bradley, M. (2020) Polymyxin-Based Photosensitizer for the Potent and Selective Killing of Gram-Negative Bacteria. Chem. Commun. 56, 3757–3760.

17. Klausen, M.; Ucuncu, M.; Bradley, M. (2020) Design of Photosensitizing Agents for Targeted Antimicrobial Photodynamic Therapy. Molecules 25 (22), 5239.

18. Digby, E. M.; Rana, R.; Nitz, M.; Beharry, A. A (2019) DNA Directed Damage Using a Brominated DAPI Derivative. Chem. Commun. 55 (67), 9971–9974.

19. Wang, Y.; Jin, Y.; Chen, W.; Wang, J.; Chen, H.; Sun, L.; Li, X. (2019) Construction of Nanomaterials with Targeting Phototherapy Properties to Inhibit Resistant Bacteria and Bio Film Infections. Chem. Eng. J. 358, 74–90.

20. Brömme, H. J.; Zühlke, L.; Silber, R. E.; Simm, A. (2008) DCFH2 Interactions with Hydroxyl Radicals and Other Oxidants - Influence of Organic Solvents. Exp. Gerontol. 43 (7), 638–644.

21. Hong, Y.; Zeng, J.; Wang, X.; Drlica, K.; Zhao, X. (2019) Post-Stress Bacterial Cell Death Mediated by Reactive Oxygen Species. Proc. Natl. Acad. Sci. U. S. A. 116 (20), 10064–10071.

22. Alves, E.; Faustino, M. A. F.; Tomé, J. P. C.; Neves, M. G. P. M. S.; Tomé, A. C.; Cavaleiro, J. A. S.; Cunha, Â.; Gomes, N. C. M.; Almeida, A. (2013) Nucleic Acid Changes during Photodynamic Inactivation of Bacteria by Cationic Porphyrins. Bioorg. Med. Chem. 21 (14), 4311–4318.

23. Boltes Cecatto, R.; Siqueira de Magalhães, L.; Fernanda Setúbal Destro Rodrigues, M.; Pavani, C.; Lino-dos-Santos-Franco, A.; Teixeira Gomes, M.; Fátima Teixeira Silva, D. (2020) Methylene Blue Mediated Antimicrobial Photodynamic Therapy in Clinical Human Studies: The State of the Art. Photodiagnosis Photodyn. Ther. 31, 101828.

24. Cieplik, F.; Pummer, A.; Regensburger, J.; Hiller, K. A.; Späth, A.; Tabenski, L.; Buchalla, W.; Maisch, T. (2015) The Impact of Absorbed Photons on Antimicrobial Photodynamic Efficacy. Front. Microbiol. 6 (JUN).

25. Tegos, G. P.; Masago, K.; Aziz, F.; Higginbotham, A.; Stermitz, F. R.; Hamblin, M. R (2008) Inhibitors of Bacterial Multidrug Efflux Pumps Potentiate Antimicrobial Photoinactivation. Antimicrob. Agents Chemother. 52 (9), 3202–3209.

26. Costa Magacho, C.; Guerra Pinto, J.; Müller Nunes Souza, B.; Correia Pereira, A. H.; Ferreira - Strixino, J. (2020) Comparison of Photodynamic Therapy with Methylene Blue Associated with Ceftriaxone in Gram-Negative Bacteria; an in Vitro Study. Photodiagnosis Photodyn. Ther. 30 (January), 101691.

27. Ortiz, M.; Fragoso, A.; Ortiz, P. J.; O’Sullivan, C. K (2011) Elucidation of the Mechanism of Single-Stranded DNA Interaction with Methylene Blue: A Spectroscopic Approach. J. Photochem. Photobiol. A Chem. 218 (1), 26–32.

28. Vardevanyan, P. O.; Antonyan, A. P.; Parsadanyan, M. A.; Shahinyan, M. A.; Hambardzumyan, L. A (2013) Mechanisms for Binding between Methylene Blue and DNA. J. Appl. Spectrosc. 80 (4), 595–599.

29. Fodor, L.; Ullmann, Y.; Elman, M. Light Tissue Interactions. In Aesthetic Applications of Intense Pulsed Light; Springer-Verlag London Limited: London; pp 11–20.

30. Xu, C.; Webb, W. W (1996) Measurement of Two-Photon Excitation Cross Sections of Molecular Fluorophores with Data from 690 to 1050 Nm. J. Opt. Soc. Am. B 13 (3), 481.

31. Trägårdh, J.; Robb, G.; Amor, R.; Amos, W. B.; Dempster, J.; McConnell, G. (2015) Exploration of the Two-Photon Excitation Spectrum of Fluorescent Dyes at Wavelengths below the Range of the Ti: Sapphire Laser. J. Microsc. 259 (3), 210–218.

32. Matsuzaki, K. (2009) Control of Cell Selectivity of Antimicrobial Peptides. Biochim. Biophys. Acta - Biomembr. 1788 (8), 1687–1692.

33. Cummings, B. S.; Wills, L. P.; Schnellmann, R. G (2004) Measurement of Cell Death in Mammalian Cells. Curr. Protoc. Pharmacol. 25 (1), 1–30.

34. Stiefel, P.; Schmidt-Emrich, S.; Maniura-Weber, K.; Ren, Q. (2015) Critical Aspects of Using Bacterial Cell Viability Assays with the Fluorophores SYTO9 and Propidium Iodide. BMC Microbiol. 15 (1), 1–9.

35. Polat, E.; Kang, K. (2021) Natural Photosensitizers in Antimicrobial Photodynamic Therapy. Biomedicines 9, 584.

36. Ragas, X.; Sanchez-Garcia, D.; Ruiz-Gonzalez, R.; Dai, T.; Agut, M.; Hamblin, M. R.; Nonell, S. (2010) Cationic Porphycenes as Potential Photosensitizers for Antimicrobial Photodynamic Therapy. J. Med. Chem. 53, 7796–7803.

37. Olson, M. E.; Ceri, H.; Morck, D. W.; Buret, A. G.; Read, R. R (2002) Biofilm Bacteria: Formation and Comparative Susceptibility to Antibiotics. Can. J. Vet. Res. 66, 86–92.

38. Kapuscinski, J. (1995) DAPI: A DNA-Specific Fluorescent Probe. Biotech. Histochem. 70 (5), 220–233.

39. Bergmann, K.; O’Konski, C. T. (1963) A Spectroscopic Study of Methylene Blue Monomer, Dimer, and Complexes with Montmorillonite. J. Phys. Chem. 67 (10), 2169–2177.

40. Demchenko, A. P (2013) Beyond Annexin V: Fluorescence Response of Cellular Membranes to Apoptosis. Cytotechnology 65 (2), 157–172.

